# Longitudinal analysis of naturally acquired antibodies to PfEMP1 CIDR domain variants and their association with malaria protection

**DOI:** 10.1101/2020.02.11.944330

**Authors:** Nyamekye Obeng-Adjei, Daniel B. Larremore, Louise Turner, Aissata Ongoiba, Shanping Li, Safiatou Doumbo, Takele B. Yazew, Kassoum Kayentao, Louis H. Miller, Boubacar Traore, Susan K. Pierce, Caroline O. Buckee, Thomas Lavstsen, Peter D. Crompton, Tuan M. Tran

**Author notes:** Both authors contributed equally (Nyamekye Obeng-Adjei, Daniel B. Larremore). Corresponding author: Tuan M. Tran.

## Abstract

Malaria pathogenicity is determined, in part, by the adherence of *Plasmodium falciparum* infected erythrocytes to the microvasculature mediated via specific interactions between PfEMP1 variant domains to host endothelial receptors. Naturally acquired antibodies against specific PfEMP1 variants can play an important role in clinical protection against malaria. We evaluated IgG responses against a repertoire of PfEMP1 CIDR domain variants to determine the rate and order of variant-specific antibody acquisition and their association with protection against febrile malaria in a prospective cohort study conducted in an area of intense, seasonal malaria transmission. Using longitudinal data, we found that IgG to the pathogenic domain variants CIDRα1.7 and CIDRα1.8 were acquired the earliest. Furthermore, IgG to CIDRγ3 was associated with reduced prospective risk of febrile malaria and recurrent malaria episodes. Future studies will need to validate these findings in other transmission settings and determine the functional activity of these naturally acquired CIDR variant-specific antibodies.

## INTRODUCTION

Malaria due to *Plasmodium falciparum* causes greater than 400,000 deaths per annum (1). Severe clinical manifestations of *P. falciparum* malaria are precipitated by widespread sequestration of infected erythrocytes (IEs) in host microvasculature including in the brain and placenta which can lead to cerebral malaria and placental malaria, respectively (2). Cytoadherence of IEs occurs via specific interactions between host endothelial receptors and *P. falciparum* erythrocyte membrane protein (PfEMP1), a parasite-derived protein expressed on the surface of IEs that is a major target of naturally acquired immunity to malaria (3-5). The PfEMP1 adhesins are encoded by ∼60 *var* gene variants that differ within and between parasite genomes and that are expressed in a mutually exclusive manner within each IE (6-8). Switching between *var* genes aids in parasite immune evasion and functional diversification of the PfEMP1 family have resulted in mutually exclusive receptor binding phenotypes correlated to differences in clinical severity (9, 10).

Members of the PfEMP1 family vary in the size and number of extracellular Duffy-binding-like (DBL) and cysteine-rich interdomain region (CIDR) domains(11). DBL and CIDR domains are classified based on sequence similarity into six (α, β, γ, δ, ε, ξ) and four (α, β, γ, δ) main classes, respectively, of which some can be further divided into sub-classes (e.g. CIDRα1.1) (12, 13). PfEMP1 generally have a semi-conserved head structure near the N-terminus consisting of a tandem DBLα-CIDR domain. This can be followed by a second DBLδ-CIDR tandem domain or additional other types of DBL domains in larger proteins. Notably, however, the VAR2CSA PfEMP1 variants do not contain typical CIDR domains and bind placental chondroitin sulfate A via specialized DBL domains (14, 15). PfEMP1 have diversified to either bind endothelial protein C receptor (EPCR) (10), the scavenger receptor CD36 (16) or yet undermined receptors via their head structure CIDR domains. These phenotypes are maintained by the chromosomal organization of the *var* genes (17). Among the subtelomeric *var* genes, Group A genes transcribed toward the telomere encode DBLα1-CIDRα1 head structures binding to EPCR or DBLα1-CIDRβ/γ/δ head structures with unknown endothelial receptor specificities. Subtelomeric Group B *var* genes transcribed toward the centromere as well as centromeric Group C *var* genes encode DBLα0-CIDRα2-6 head-structures binding to CD36. In addition to this, chimeric group B/A *var* genes encode EPCR-binding DBLα0-CIDRα1 head structures. The EPCR-binding phenotype has been implicated in severe malaria (18-21), whereas CD36 binding has been associated with uncomplicated malaria (22, 23). Severe malaria has been associated with rosetting, a phenomenon which involves binding between an IE and several uninfected erythrocytes but with unclear clinical significance. A set of group A PfEMP1 with DBLα1-CIDRβ/γ/δ domains have been shown to mediate rosettes.

Immunity to severe malaria is generally acquired after only one to two severe episodes (24) with naturally acquired antibodies specific for PfEMP1 variants likely playing an important role in clinical protection (25). Antibodies to group A PfEMP1 variants tend to be acquired prior to antibodies to group B and C variants (26) and are associated with protection from severe malaria (27). Similarly, antibodies to EPCR-binding CIDRα1 domains are acquired more rapidly than antibodies to other CIDR domains in areas of high malaria transmission intensity and are boosted by severe malaria but not uncomplicated malaria (28, 29). However, a recent study showed that antibodies to both rosetting-associated DBLα variants and CD36-binding CIDR domains predicted reduced risk of severe malaria to a similar extent as antibodies to EPCR-binding CIDR domains (30). The same study also showed that antibodies to group 2 DBLα variants, which are associated with rosetting (31), also predicted protection from uncomplicated malaria.

To gain further insight into the role of PfEMP1-variant specific antibodies, we assessed IgG responses against a repertoire of PfEMP1 CIDR domains to determine the rate and order of variant-specific antibody acquisition and their association with protection against uncomplicated febrile malaria in a prospective cohort study conducted in a Malian village with intense and seasonal malaria transmission.

## RESULTS

### IgG specific for CIDRα1, CIDRδ, and CIDRγ domain variants are acquired rapidly

Naturally acquired IgG antibody responses to 35 PfEMP1 CIDR domain variants representing subtypes α, γ and δ CIDR, as well as three well-studied *P. falciparum* antigens (circumsporozoite protein [PfCSP], apical membrane protein 1 [PfAMA1], and merozoite surface protein 1 [PfMSP1]), tetanus toxoid (non-malaria positive control), and bovine serum albumin (non-specific background control; Table S1) were determined by multiplex bead-based immunoassay in 680 children and adults from the Kalifabougou, Mali cohort at their healthy baseline in May 2011 (Fig. S1). Hierarchical clustering of baseline PfEMP1-specific IgG reactivity revealed distinct clustering of samples by age, and by the presence of PCR-documented, asymptomatic *P. falciparum* infection; as well as clustering of antigen targets by group (A, B, or B/A), binding phenotype (EPCR, CD36, or unknown), and CIDR domain class (Fig. 1a); suggesting differential rates of acquisition of IgG between PfEMP1 variants with cumulative *P. falciparum* exposure and the acquisition of clinical immunity to malaria. PfEMP1-specific IgG reactivity increased rapidly up to 8 years of age, and within each age stratum, *P. falciparum* PCR-positive individuals exhibited greater variant-specific IgG reactivity than uninfected individuals (Fig. 1b).

**Fig. 1.**
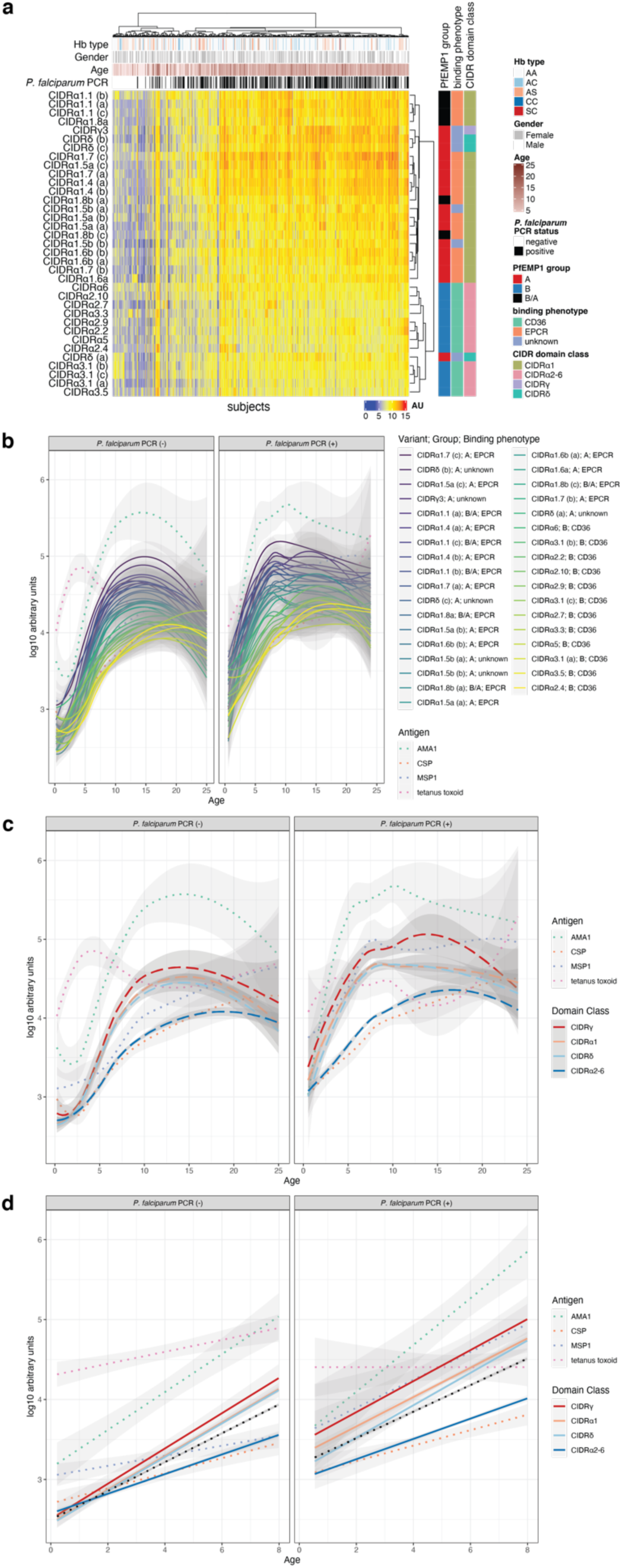
IgG antibodies to PfEMP-1 variants belonging to the A or B/A groups or having the EPCR-binding phenotype are rapidly acquired during childhood. **a** Hierarchical clustering heatmap showing IgG reactivity to each of the 35 PfEMP-1 variants in 680 subjects at enrollment (May 2011 healthy baseline). Clustering was performed using the Ward.D method and the Pearson distance metric. AU refers to arbitrary concentration units, which was calculated by fitting data to a dilutional standard curve of pooled hyper-immune plasma from malaria-exposed Malian adults. **b-c** IgG reactivity obtained at May 2011 healthy baseline versus age for each PfEMP-1 variant (solid lines) or grouped by CIDR domain class (dashed lines) with loess fit curves and 95% confidence intervals. Control antigens shown as dotted colored lines. **d** Linear portion of plot in **b** (age range 3 months to 8 years) with linear fit curves and 95% confidence intervals (see Table S2). For comparison, regression line for all variants together is represented by the black dotted line.

Categorization of PfEMP-1 variants by CIDR domain class suggested that IgG specific for variants in the CIDRγ, CIDRα1, and CIDRδ classes was acquired rapidly whereas IgG specific for group B variants of the CIDRα2-6 class was acquired slowly irrespective of *P. falciparum* infection status (Fig. 1c). Indeed, when compared to variants of other domain classes within the linear range of the fit curves (<8 years of age), IgG specific for variants within each of the CIDRγ, CIDRα1, and CIDRδ classes increased significantly more rapidly with age, whereas IgG specific for variants of the CIDRα2-6 classes increased significantly more slowly with age, independent of *P. falciparum* infection status (Fig. 1d and Table S2). Of note, IgG specific for AMA1, CSP, and MSP1 increased predictably with age in early childhood and plateaued in adolescence or young adulthood, which is similar to what we previously observed in this cohort (32, 33) (Fig. 1b-c). As we observed previously by ELISA in a separate cohort in Mali (34), increases in tetanus toxoid-specific IgG in early childhood and adolescence corresponded with the primary childhood vaccine series (diphtheria, tetanus, pertussis) and a subsequent booster of a tetanus toxoid-containing vaccine in females of child-bearing age (Fig. 1b-c).

With the exclusion of children <6 months of age whose IgG is most likely maternally derived, ranking of antigens by decreasing seropositivity within each age group revealed immunodominance of CIDRα1, CIDRδ, and CIDRγ domain classes, which are all either of the A or B/A *var* group, in early childhood (<7 years) that is maintained to a large degree in adolescence and early adulthood (Fig. S2). Notably, the most prevalent PfEMP1-specific IgG reactivity among individuals greater than 1 year of age was against CIDRα1.7(c) with seroprevalence rapidly rising from 25% in 2 to 3-year-old children to 60% in 4 to 6-year-old children and surpassing 95% in older children and adults (Fig. S2). However, the majority of individuals within the oldest age group (15-25 years) were also seropositive for several variants within the CIDRα2-6 domain classes, suggesting that IgG antibodies against these variants are eventually acquired with additional years of malaria exposure.

To assess the longitudinal acquisition of variant-specific IgG, we determined variant-specific IgG reactivity across five annual cross-sectional surveys conducted just prior to each malaria transmission season for an age-stratified random sample of 60 children from the entire cohort (Fig. S1). Children in this subset experienced a median of 6 febrile malaria episodes (interquartile range, 4−9 episodes) with a broad range of parasite densities and distributed widely but with clear seasonal peaks in the number of episodes during the five-year surveillance period (Fig. 2a-b). In the youngest children (6 months to 2 years), IgG specific for variants of the CIDRα1 and CIDRδ domain classes began low and then increased rapidly over four malaria seasons, whereas IgG specific for CIDRγ initially decreased during the first two years before rising during the third year of surveillance (Fig. 2c). In contrast, older children (3 to 8 years) appeared to maintain stable levels of IgG specific for all PfEMP1 variants over four malaria seasons (Fig. 2c).

**Fig. 2.**
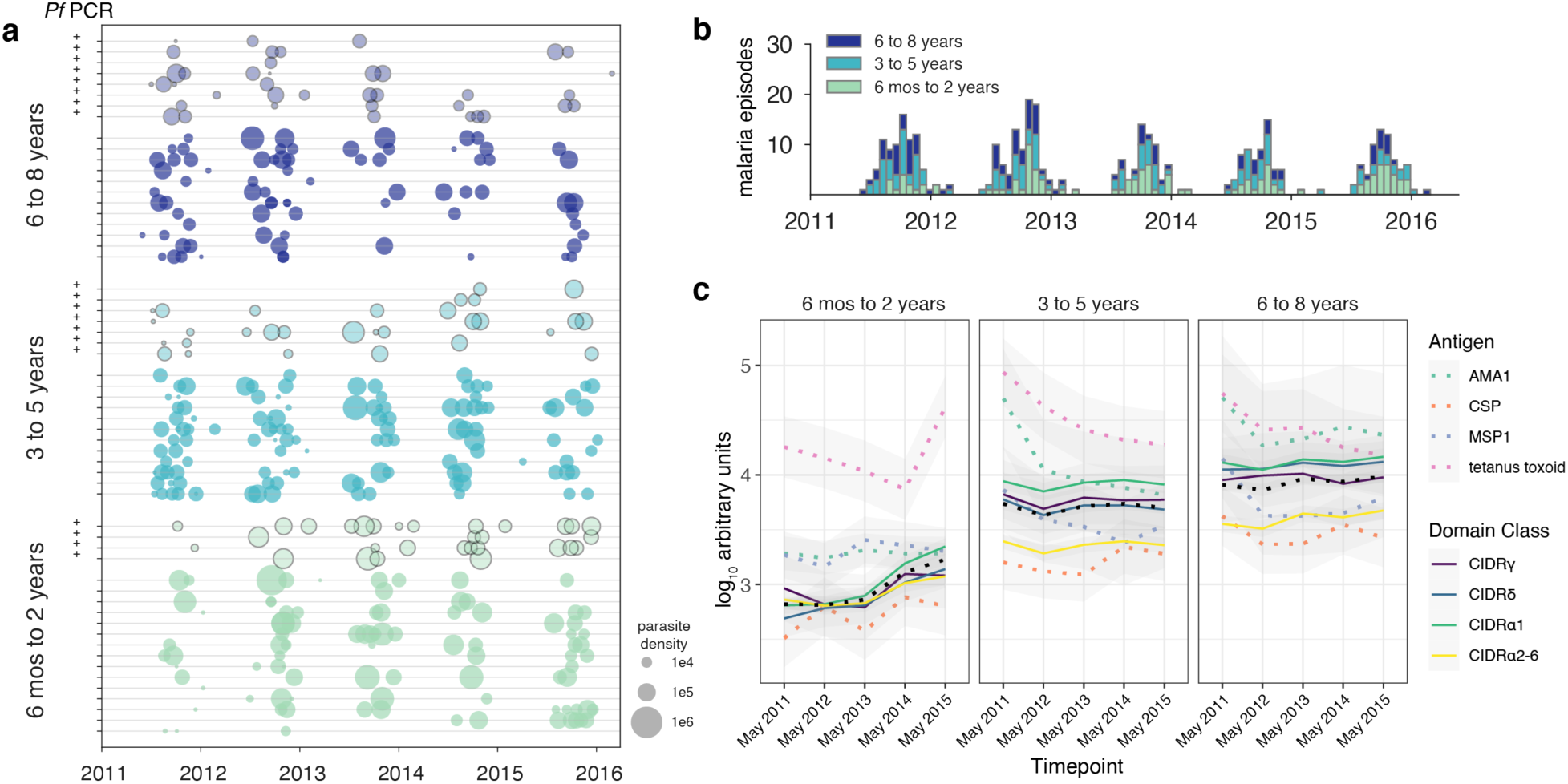
Longitudinal analysis of PfEMP-1 variant-specific IgG over multiple malaria seasons. IgG reactivity specific to PfEMP-1 variants was determined for 60 children ages 6 months to 8 years (**see Figure S1**) at five cross-sectional surveys prior to the malaria season. **a** Malaria incidence over five malaria seasons for 60 children in children aged 6 months to 2 years; 3 to 5 years; and 6 to 8 years (n = 20 per age group). Plus (+) signs in the left margin indicate subjects with asymptomatic *P. falciparum* parasitemia at enrollment. Size of bubble is proportional to parasite density determined at each visit. **b** Number of malaria episodes per two-week period by age group over five malaria seasons. **c** Longitudinal IgG reactivity at five cross-sectional surveys in the same children. Color scale for variant is ordered by slopes estimated from cross-sectional data (**Table S2**) to facilitate comparison.

### Acquisition of IgG antibodies to CIDR domain classes is highly ordered with IgG against EPCR-binding domain variants CIDRα1.7 and CIDRα1.8 acquired first

We next asked whether IgG antibodies to individual PfEMP1 variants were acquired in a particular order. Here we used an approach called *minimum violations ranking* (MVR), where an algorithm searches over different possible orders of acquisition of antibodies to PfEMP1 variants such that, if a particular an order is assumed for each child, the number of order violations observed in the data overall is minimized (refer to methods). We observed significantly less violations if we assumed an ordered acquisition of antibodies compared to a model with randomized seroconversion orders for each child, which highly suggests a hierarchical exposure to different parasite CIDR domains in this population (Fig. 3a-d). At the variant level, IgG specific to CIDRα1.7(c) was acquired first followed by IgG to CIDRα1.8b(a), CIDRα1.8b(c), CIDRα1.7(a), CIDRα6, and CIDRγ3 (Fig. 3a). Grouped by CIDR domain class, IgG was acquired against CIDRγ first followed by CIDRα1, CIDRδ, and CIDRα2-6 (Fig. 3b). Grouped on the basis of upstream sequence, IgG was acquired against B/A first followed by A and B (Fig. 3c). Lastly, when variants were grouped by binding phenotype, IgG against EPCR-binding domains were acquired first followed by domains with unknown binding phenotypes and CD36-binding domains (Fig. 3d). Whether this reflects differential prevalence of variants in the parasite population or age-specific expression patterns remains an open question.

**Fig. 3.**
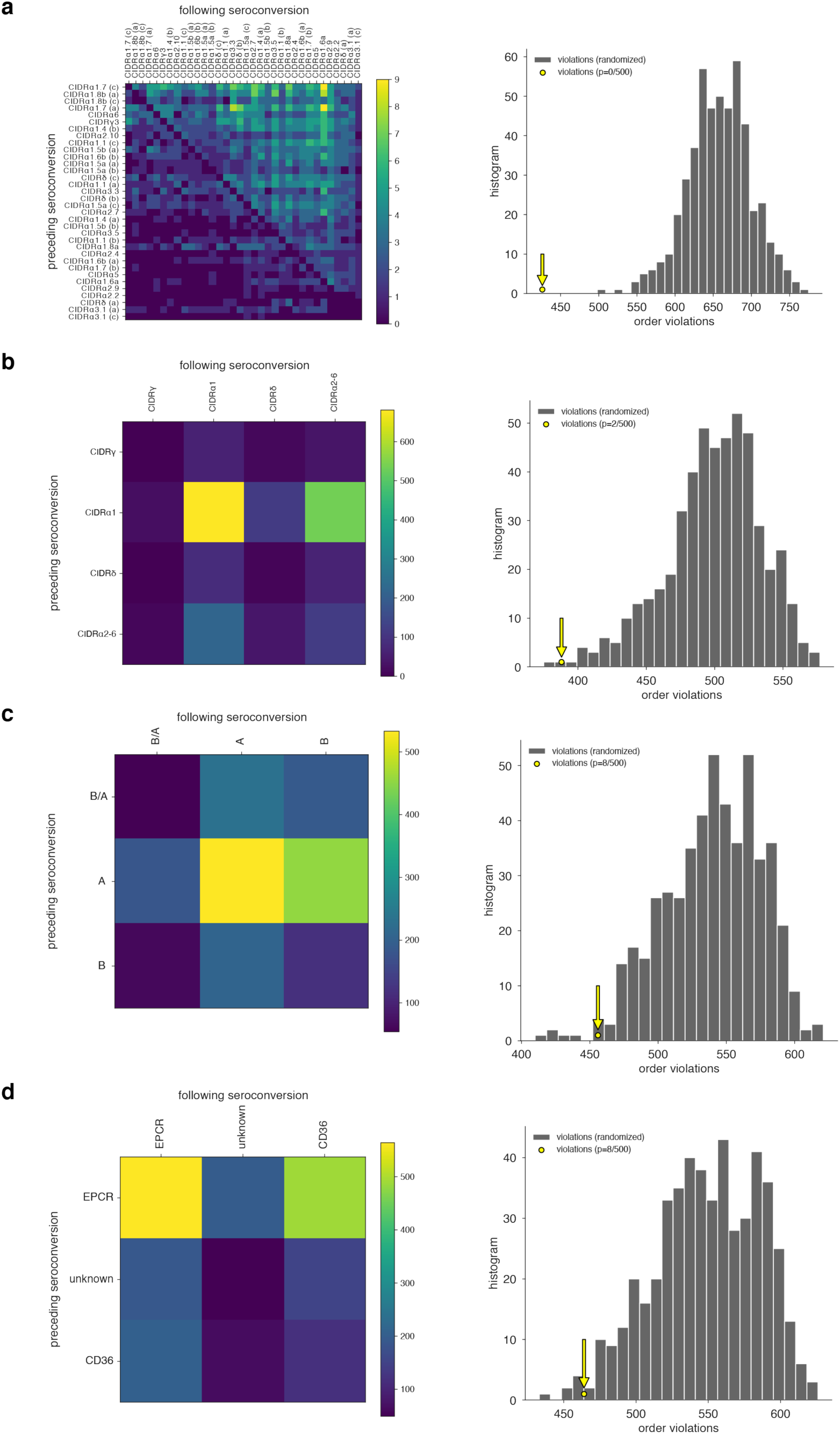
Acquisition of IgG to PfEMP-1 variants over time is hierarchical. Using longitudinal data, seropositivity was determined for each variant within each subject at each time point to determine the year of seroconversion. Seroconversion year was then used to generate a matrix representing the number of times that seroconversion for a variant (rows) precedes another variant (column) across all subjects. To find consensus ordering, the matrix was sorted to minimize the number of violations. Observed consensus ordering was compared against 200 independent procedures in which the seroconversion orders for each subject was randomized and consensus ordering was carry out in the same manner (right panel). Analysis was performed at the level of **a** individual variants, **b** CIDR domain class, **c** upstream sequence group, and **d** binding phenotype.

### CIDRγ-specific IgG associates with protection from uncomplicated, febrile malaria

We focused on the risk of uncomplicated malaria given that severe malaria was rarely observed in the Kalifabougou cohort due to early diagnosis and treatment. We specifically evaluated whether baseline seropositivity for each variant could predict protection from febrile malaria after subsequent PCR-confirmed *P. falciparum* parasitemia in individuals who began the study PCR-negative using a Cox regression model that included age, presence of the malaria-protective HbS allele, gender, IgG reactivity to AMA1 (as a surrogate for prior malaria exposure), and seropositivity to each of the 35 PfEMP-1 variants as covariates. Notably, seropositivity to CIDRγ3 (IT4var08), which has an unknown binding phenotype was significantly associated with reduced risk of febrile malaria (Table 1). CIDRγ domains have been associated with rosetting of erythrocytes (11), a phenomenon associated with severe forms of malaria (35) except in individuals with blood group O erythrocytes which appear to exhibit reduced rosetting (36). We therefore hypothesized that the reduced risk afforded by CIDRγ-specific IgG might occur via the inhibition of rosette formation and may therefore be negatively affected by blood group O. When included as a covariate in a reduced Cox regression model, group O blood type affected neither malaria risk itself nor the association between CIDRγ-specific IgG and risk of febrile malaria (Table S3). Notably, baseline CIDRγ3-specific IgG reactivity did not significantly correlate with decreased parasite density at the first malaria episode after controlling for age and the presence of the HbS allele (data not shown), suggesting that CIDRγ-specific IgG may not have anti-parasite activity. Given the association between CIDRγ-specific IgG and delay in malaria fever during the first year of the study, we specifically examined if CIDRγ3 serostatus at the beginning of each malaria season affected the risk of recurrent malaria episodes in the 60 children who were longitudinal evaluated for PfEMP1 IgG responses over five malaria seasons. Presence of CIDRγ3-specific IgG prior to each season predicted a reduction in febrile malaria episodes even after controlling for AMA1-specific IgG serostatus and the HbS allele (Table 2).

**Table 1.** Relationship between CIDR variant seropositivity and protection from febrile malaria. Results of Cox regression model assessing PfEMP1 variant-specific IgG on the risk of febrile malaria after incident *P. falciparum* infection in which covariates were age, gender, presence of the HbS allele, and IgG seropositivity for five CIDR variants selected using the least absolute shrinkage and selection operator (LASSO; refer to Methods). Analysis was restricted to subjects who were at least 6 months of age and began the study negative for *P. falciparum* infection by PCR (271 subjects). Malaria risk was determined based on time to clinical malaria, defined as axillary temperature >37.5 degrees C and any parasitemia, after PCR-documented blood-stage infection (163 malaria events). Follow-up time was limited to 60 days from initial blood-stage infection. Results are ordered by increasing P values. HR = hazard ratio; LCI = lower 95% confidence interval; UCI = upper 95% confidence interval.

| Variant | Number seropositive (N=271) | Alternate name | PfEMP1 group | Binding Phenotype | Genome/ Isolate | HR | LCI | UCI | P value |
| --- | --- | --- | --- | --- | --- | --- | --- | --- | --- |
| CIDRγ3 | 129 | IT4var08 | A | unknown | IT4 | 0.607 | 0.42 | 0.876 | 0.00763 |
| CIDRα3.1 (a) | 17 | DD2var01 | B | CD36 | DD2 | 0.208 | 0.0523 | 0.828 | 0.0259 |
| CIDRα3.3 | 55 | IT4var26 | B | CD36 | IT4 | 0.806 | 0.458 | 1.42 | 0.454 |
| CIDRα2.10 | 37 | IT4var30 | B | CD36 | IT4 | 0.861 | 0.507 | 1.46 | 0.58 |
| CIDRα2.9 | 25 | IT4var45 | B | CD36 | IT4 | 0.83 | 0.429 | 1.6 | 0.58 |

**Table 2.**
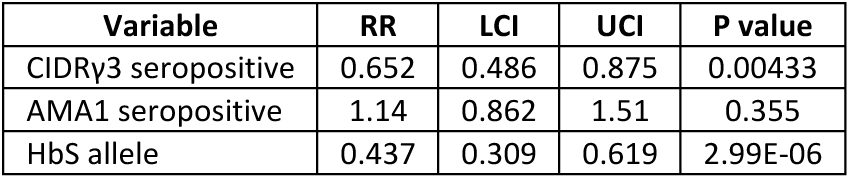
Relationship between CIDRγ3 seropositivity and protection from recurrent malaria episodes. Results of the Andersen-Gill extension of the Cox regression model to assess the relationship between CIDRγ3-specific IgG seropositivity and the risk of recurrent febrile malaria episodes (defined as fever >37.5°C and any parasitemia; 376 events) in 60 children who were followed longitudinally over five malaria transmission seasons from 2011 through 2015. Presence of the HbS allele and AMA1 seropositivity, a surrogate for overall malaria exposure, were included as covariates. CIDRγ3-specific and AMA1-specific IgG seropositivity were treated as time-dependent covariates that varied over each season. RR = relative risk; LCI = lower 95% confidence interval; UCI = upper 95% confidence interval.

## DISCUSSION

PfEMP1 variants containing domains of the CIDRα1 class generally bind to EPCR on endothelial cells and are associated with severe malaria (10), whereas variants containing domains of the CIDRα2-6 classes bind to CD36 present on several host cell types, including microvascular endothelial cells, mononuclear phagocytes, and platelets (16, 37). Antibodies targeting these PfEMP1 domains can potentially disrupt adhesion of IEs to host receptors but can also facilitate IE clearance via opsonization and phagocytosis or antibody-mediated cytotoxicity (10, 38, 39). Consistent with a prior study conducted in a Tanzanian cohort (28), we observed early acquisition of IgG antibodies against EPCR-binding PfEMP1 variants of the CIDRα1 domain class relative to CD36-binding variants in both age-stratified cross-sectional and longitudinal analyses. This is also consistent with studies that investigated acquisition of antibodies to PfEMP1 classified by upstream sequence group and found that antibodies to DBL and CIDR domains belonging to group A and B/A are acquired earlier in life than group B and C variants among individuals living in malaria-endemic settings (26, 40). Importantly, the antigen panel used in the current study contained unique CIDR domains not covered by these prior studies.

Among the 35 distinct CIDR domains evaluated here, CIDRα1.7(c) elicited the most robust and prevalent IgG responses in early childhood, eventually approaching 100% seroprevalence in adolescents and adults in this cohort. Longitudinal analysis to assess hierarchical acquisition confirmed that IgG antibodies specific for CIDRα1.7(c) were acquired first, with IgG against the related CIDRα1.7(a) variant acquired fourth. Transcripts encoding CIDRα1.7 domains have been found to predominate among the most severe cases of pediatric cerebral malaria—those that lead to brain swelling and death (19). The immunodominance of CIDRα1.7(c) may be a consequence of epitopes targeted by cross-reactive CIDRα1 antibodies (41, 42). Moreover, PfEMP1 with CIDRα1.4 and CIDRα1.7 domains frequently contain ICAM1-binding DBLβ domains (43). The dual EPCR- and ICAM1-binding phenotype is thought to be particularly pathogenic, and antibodies to these DBLβ domains have been associated with reduced risk of clinical malaria with parasite densities of ≥10,000 parasites/μl (44). We also observed early acquisition of IgG specific for CIDRα1.8 domains. Expression of these domains, as well as EPCR-binding CIDRα1 domains in general, is associated with severe malaria including cerebral malaria in African children (18-21, 45) and Indian adults (46).

Given that all CIDRα1 variants have been linked to severe malaria in African children, the early acquisition of IgG specific to CIDRα1.7 and CIDRα1.8 domains may just be a reflection of local parasite population dynamics rather than enhanced pathogenicity conferred by these specific CIDR variants. However, the potential lethality of parasites expressing CIDRα1 in general underscores why a vigorous host antibody response against these variant domains in early childhood may be advantageous. This study builds on older work (4, 47, 48) showing an age-specific acquisition of antibodies to particular parasite strains, and we are able to statistically confirm this pattern for the first time, and identify key genetic underpinnings of those observations. We still cannot address the slippery problem of whether this order reflects the circulation of genotypes with different transmissibility; under this scenario, high fitness genotypes lead to high prevalence and therefore low age of first infection, and coincidentally cause more disease in relatively non-immune children compared to low fitness genotypes as a result. In contrast, it is possible that the ordered expression of PfEMP1 variants across strains, potentially in response to the immune status of the parasite’s immediate host, leads to the hierarchical acquisition of antibodies observed.

Due to the low incidence of severe disease in the cohort, we could not assess the impact of CIDRα1.7-specific or CIDRα1.8-specific antibodies on the risk of severe malaria in the study. However, when all 35 CIDRs were assessed for association with the prospective risk of uncomplicated, febrile malaria, IgG specific to CIDRγ3 (IT4var08) was significantly associated with reduced malaria risk. PfEMP1 variants encoding CIDRβ, CIDRγ, or CIDRδ domains have been associated with rosetting (11, 46), which can enhance microvasculature obstruction thereby increasing malaria severity. However, direct evidence that any of these CIDR domains have intrinsic rosetting properties is lacking (49). Rather, their association with rosetting may be related to their tandem expression with an adjacent DBLα1 at the N-terminal head (50). Rosetting frequency has been correlated with severity of malaria with the highest levels in cerebral malaria (35, 51, 52) but is still commonly observed in uncomplicated malaria. Thus, the role of rosetting in severity of malarial disease remains unclear. Nevertheless, disruption of rosettes by targeting DBL1α has been used as a vaccine strategy (53), and antibodies to rosetting-associated group 2 DBLα domains predicted protection from uncomplicated malaria, suggesting a protective role for these antibodies in less severe disease (30, 31). Although speculative, it is possible that naturally acquired CIDRγ-specific IgG confers protection from febrile malaria by blocking rosette formation. However, this mechanism is not supported by the current study given that the protection attributable to CIDRγ-specific IgG is unchanged after controlling for blood group O, which has been shown to be protective against severe falciparum malaria through the reduction of rosetting (36). It also must be noted that reduced malaria risk was not observed for IgG-specific for variants of the CIDRδ class, which is also predicted to have rosetting activity. Furthermore, as CIDRγ3 was the only CIDRγ domain variant tested in this study, it remains unknown whether the protective effect observed here would be generalizable to IgG targeting other CIDRγ variants.

A limitation of the study is that we did not sequence *var* transcripts from individuals with *P. falciparum* infections in the longitudinal analysis. This may have allowed us to prospectively assess if seroconversion against specific CIDRs such as CIDRα1.7, CIDRα1.8, or CIDRγ reliably led to the absence of parasites expressing the corresponding *var* transcript during clinical malaria episodes. In addition to our limited assessment of CIDRγ domains, we also did not evaluate CIDRβ domains, which also have been associated with the rosetting phenotype.

In summary, this longitudinal study provides evidence that acquisition of IgG antibodies to PfEMP1 variants is ordered and demonstrates that antibodies to CIDRα1 domains, specifically the pathogenic domain variants CIDRα1.7 and CIDRα1.8, are acquired the earliest in children residing in an area of intense, seasonal malaria transmission. We also show that IgG antibodies to the rosetting-associated CIDRγ3 domain is acquired early and is associated with protection from febrile malaria. Future studies will need to validate these findings in other transmission settings and determine the functional activity of these naturally acquired CIDR variant-specific antibodies.

## METHODS

### Ethics

The Ethics Committee of the Faculty of Medicine, Pharmacy and Dentistry at the University of Sciences, Techniques, and Technology of Bamako, and the Institutional Review Board of the National Institute of Allergy and Infectious Diseases, National Institutes of Health approved this study (ClinicalTrials.gov identifier: NCT01322581). Written, informed consent was obtained from the parents or guardians of participating children or from adult participants.

### Study Site

The study was conducted in the village of Kalifabougou, Mali, which is located 40 km northwest of Bamako, Mali within the savanna ecoclimatic zone. Within this community, Bambara is the predominant ethnic group, and ∼90% of residents engage in subsistence farming. Malaria transmission is intense and seasonal, reliably occurring from June through December, with the vast majority of malaria cases caused by *P. falciparum* (54).

### Study Population and Study Design

Recruitment and enrollment procedures of participants for this study have been previously described (55). Briefly, exclusion criteria at enrollment included a hemoglobin level <7 g/dL, axillary temperature ≥37.5°C, acute systemic illness, underlying chronic disease, use of antimalarial or immunosuppressive medications in the past 30 days, or pregnancy. The study design and selection of subjects are summarized in Fig. S1.

### Human samples

At the beginning and end of the malaria-transmission season, blood samples were drawn by venipuncture into sodium-citrate-containing Vacutainer tubes (Becton Dickinson). Plasma was separated by centrifugation and cryopreserved. Hemoglobin typing was performed using a D-10 instrument (Bio-Rad). Blood for ABO typing was collected in EDTA containing microtainers. ABO typing was conducted with forward typing using Cypress Diagnostics Reagents. Anti-A, Anti-B, and Anti-AB IgM reagents were mixed with the sample, and blood type was determined by agglutination. During the first malaria season, blood was collected by finger-prick onto 903 filter paper (Whatman) for PCR analysis at each scheduled clinic visit (occurring at 2-week intervals for 7 months) and sick visit for subsequent molecular diagnostics.

### Diagnosis and Treatment of Infections

#### Clinical malaria episodes

Individuals were initially enrolled in May 2011 and have been followed continuously since unless withdrawn or lost to follow-up. During the first malaria season, clinical malaria episodes were detected prospectively by self-referral and weekly active clinical surveillance visits which alternated between the study clinic and the participants’ homes. Passive malaria surveillance and pre- and post-malaria season cross-sectional surveys have continued during subsequent years. All individuals with signs and symptoms of malaria and any level of *Plasmodium* parasitemia detected by light microscopy were treated according to the National Malaria Control Program guidelines in Mali. For the current study, a clinical malaria episode was defined as any parasitemia on contemporaneous blood smear, an axillary temperature of ≥37.5°C within 24 hours, and no other cause of fever discernible by physical exam.

#### Blood smears

Thick blood smears were stained with Giemsa and counted against 300 leukocytes. Parasite densities were recorded as the number of asexual parasites/µl of blood based on a mean leukocyte count of 7500 cells/µl. Each smear was read in blinded manner by two certified microscopists of the laboratory team.

#### Molecular detection

For each participant, the first *P. falciparum* infection of the initial malaria season was detected retrospectively by PCR analysis of the longitudinally collected dried blood spots (54). First malaria episodes were determined from the clinical visit data.

### Protein Expression and Multiplex Immunoassays

The 35 recombinant His-tagged CIDR domains (Table S1) were expressed in baculovirus-transfected insect cells, and purified by nickel affinity chromatography as previously described (28, 42, 56). AMA1, CSP, and MSP1 recombinant proteins were kindly provided by David Narum (Laboratory of Malaria Immunology and Vaccinology, NIAID, NIH). AMA1 and CSP were expressed from *P. falciparum* 3D7 in *P. pastoris* as previously described (57, 58). MSP1 was expressed from *P. falciparum* 3D7 in *Escherichia coli* as previously described (59). Purified tetanus toxoid was provided the staff at Biologic Laboratories, University of Massachusetts Medical School. Bovine serum albumin (BSA) was obtained from Sigma. These proteins were coupled to MagPlex-C micospheres (Luminex) and mixed to form a protein bead array in which IgG reactivity to each antigen could be measured in multiplex, as previously described(60) with minor modifications. Briefly, plasma samples were diluted 1:500 and 1:2000 (to better assess highly reactive antigens) in Assay Buffer E (ABE: 0.1% BSA, 0.05% Tween-20 in PBS, pH7.4). For each plate, pooled malaria-hyperimmune plasma was serially diluted in ABE at 1:50, 1:158, 1:500, 1:1580, 1:5000, 1:1580, 1:50000, and 1:158000 to generate an 8-point dilutional standard curve. 50 µl of beads and 50 µl of diluted plasma was added to 96-well microtiter plates (MSBVS 1210, Millipore, USA) pre-wetted with ABE. 50 µl of phycoerythrin-conjugated Goat Anti-Human IgG (Jackson ImmunoResearch Laboratories), diluted 1:3000 was added, and mean fluorescent intensities were measured using the Luminex 200 system. To account for plate-to-plate variation, fluorescence intensities were normalized using the median reactivity for each antigen on each plate. Normalized intensities were then scaled to the mean reactivity for each antigen to allow comparison between antigens. Using the ncal function within the nCal package(61), IgG concentrations were interpolated from the standard curves generated from serial diluted pooled malaria-immune plasma and the resulting concentrations reported as arbitrary units (AU), which was used for statistical analyses and visualization.

### Ordered Acquisition Analysis

If seroconversion to CIDR domains occurs in a stereotypical order, then each individual’s sequence of seroconversions in this longitudinal study should be congruent with that order. Of course, we do not know such an order *a priori*, so we find it by searching over all orderings to find the one that minimizes the number of order-violating seroconversions. This *minimum violations ranking* (MVR) consists of both the ordinal ranking itself and a corresponding number of rank violations *v*. These outputs can be visualized by plotting a heatmap, with indices ordered by the minimizing ranking as in Fig. 3A. Clear triangular structure indicates the strength of the ordering, and *v* is equal to the sum of the sparser triangle.

Note that the more that individuals’ seroconversions occur strictly in their rank order, the smaller *v* will be. In this way, the number of violations *v* provides a convenient test statistic for a standard one-tailed p-value test: our null hypothesis is that there is no meaningful order to seroconversions, and thus, randomly permuting the order of seroconversions for each individual and recomputing *v* should make no difference. In other words, the null hypothesis is that the number of violations *v* in the real data is statistically indistinguishable from the number of violations in the time-randomized data *v*_random_. The p-value can be computed then as p=Pr(*v* < *v*_random_). When actual seroconversions are significantly more orderable than random seroconversions (while preserving the seroconversion counts per individual and seroconversions per CIDR domain), it indicates the presence of a statistically significant stereotypical ordering, as in Fig. 3B.

Computations were performed according to the following details. Let matrix entry A_ij_ be the number of times, over each individual, that a seroconversion to i was observed prior to a seroconversion to j. If the matrix’s rows and columns are sorted according to some re-ordering r, then the number of violations v can be computed as the sum of the lower triangle of A(r). Finding the r that minimizes v can be done by beginning from a random r, and then sequentially proposing swaps of pairs of indices in which any swap that increases v is rejected and otherwise swaps are accepted. This MVR algorithm exits after a large number of proposed swaps have been rejected without any decrease in v, and the output is both v and the order of seroconversion that corresponds to that v. Permutation tests were then performed by shuffling the seroconversions and years, independently for each individual, and then applying the computation above.

## Statistical Analysis

The use of specific statistical tests and methods are indicated in the Results and/or figure legends. Statistical significance was defined as a 2-tailed P value of <.05. Analyses were performed in R version 3.6.1 (http://www.R-project.org). Plots were generated with the *ggplot2* package. Cox regression was performed using the *survival* and *glmnet* packages. For the time to febrile malaria analysis (Table 1), variable selection from among the 35 CIDR seropositivity variables, age, gender, AMA1 seroreactivity, and the presence of the HbS allele was determined using regularized Cox regression fit with the least absolute shrinkage and selection operator (LASSO) penalty using 10-fold cross validation with 1000 iterations (62). The follow-up period after initial blood-stage infection was 60 days. For the final Cox regression model, age, gender, and the presence of the HbS allele were included as co-variates along with the LASSO-selected CIDR variables. For the recurrent event analysis, the Andersen-Gill extension of the Cox regression model was used to determine the relative risk of malaria over five malaria seasons using presence of the HbS allele as a covariate and AMA1-specific IgG seropositivity (a surrogate for overall malaria exposure) and CIDRγ3-specific IgG seropositivity as time-dependent covariates that varied over each season.

## Acknowledgments

We thank all the participants in the Kalifabougou cohort and the field team for making this study possible. We also thank David Narum (Laboratory of Malaria Immunology and Vaccinology, NIAID, NIH) for providing the recombinant AMA1, CSP, and MSP1 proteins and Chiung-Yu Huang (University of California, San Francisco) for her critical review and suggestions regarding the survival and recurrent event analysis. The staff at Biologic Laboratories, University of Massachusetts Medical School at Jamaica Plains, MA generously provided the purified tetanus toxoid.

## Author Contributions

NO, LHM, SKP, LT, TL, PDC, and TMT conceived the study. AO, SD, KK, and BT were responsible for the cohort study and collection of samples. NO, LT, SL, and TBY conducted the experiments. NO, DBL, and TMT analyzed the data. TMT, TL, CB, DBL, and PDC wrote the manuscript with contributions from NO, LT, LHM, and SKP. All authors read and approved the manuscript.

