## Supplemental Information for "Longitudinal analysis of naturally acquired antibodies to PfEMP1 CIDR domain variants and their association with malaria protection"

**Table S1. Antigens tested in multiplex immunoassay**.

| Original Antigen Labels | Figure Labels | Group | Binding Phenotype | Genome/ Isolate | Domain Class | Name used in figures of Rambathla et al PMID. 30365003 |
|---|---|---|---|---|---|---|
| HB3var03 | CIDRα1.4 (a) | A | EPCR | HB3 | CIDRα1 | α1.4 (a) |
| IT4var7 | CIDRα1.4 (b) | A | EPCR | IT4 | CIDRα1 | α1.4 (b) |
| X1965_2 | CIDRα1.5a (a) | A | EPCR | 1965 | CIDRα1 | α1.5a (a) |
| ERS010323 | CIDRα1.5a (b) | A | EPCR | GA013 | CIDRα1 | α1.5a (b) |
| ERS01002 | CIDRα1.5a (c) | A | EPCR | GA014 | CIDRα1 | α1.5a (c) |
| X198_5 | CIDRα1.5b (a) | A | unknown | 1918 | CIDRα1 | α1.5b (a) |
| X198313 | CIDRα1.5b (b) | A | unknown | 1983 | CIDRα1 | α1.5b (b) |
| HB3var02 | CIDRα1.6a | A | EPCR | HB3 | CIDRα1 | α1.6a |
| ERS01057 | CIDRα1.6b (a) | A | EPCR | GA018 | CIDRα1 | α1.6b (a) |
| ERS01003 | CIDRα1.6b (b) | A | EPCR | GA019 | CIDRα1 | α1.6b (b) |
| X1965_8 | CIDRα1.7 (a) | A | EPCR | 1965 | CIDRα1 | α1.7 (a) |
| X1918_3 | CIDRα1.7 (b) | A | EPCR | 1918 | CIDRα1 | α1.7 (b) |
| ERS01043 | CIDRα1.7 (c) | A | EPCR | GA024 | CIDRα1 | α1.7 (c) |
| IT4var08 | CIDRγ3 | A | unknown | IT4 | CIDRγ | γ |
| HB3var05 | CIDRδ (a) | A | unknown | HB3 | CIDRδ | δ (a) |
| HB3var35 | CIDRδ (b) | A | unknown | HB3 | CIDRδ | δ (b) |
| IT4var02 | CIDRδ (c) | A | unknown | IT4 | CIDRδ | δ (c) |
| IT4var30 | CIDRα2.10 | B | CD36 | IT4 | CIDRα2-6 | α2.10 |
| IT4var24 | CIDRα2.2 | B | CD36 | IT4 | CIDRα2-6 | α2.2 |
| IT4var33 | CIDRα2.4 | B | CD36 | IT4 | CIDRα2-6 | α2.4 |
| IT4var61 | CIDRα2.7 | B | CD36 | IT4 | CIDRα2-6 | α2.7 |
| IT4var45 | CIDRα2.9 | B | CD36 | IT4 | CIDRα2-6 | α2.9 |
| DD2var01 | CIDRα3.1 (a) | B | CD36 | DD2 | CIDRα2-6 | α3.1 (a) |
| HB3var27 | CIDRα3.1 (b) | B | CD36 | HB3 | CIDRα2-6 | α3.1 (b) |
| IT4var21 | CIDRα3.1 (c) | B | CD36 | IT4 | CIDRα2-6 | α3.1 (c) |
| IT4var26 | CIDRα3.3 | B | CD36 | IT4 | CIDRα2-6 | α3.3 |
| IT4var15 | CIDRα3.5 | B | CD36 | IT4 | CIDRα2-6 | α3.5 |
| IT4var14 | CIDRα5 | B | CD36 | IT4 | CIDRα2-6 | α5 |
| IT4var12 | CIDRα6 | B | CD36 | IT4 | CIDRα2-6 | α6 |
| IT4var20 | CIDRα1.1 (a) | B/A | EPCR | IT4 | CIDRα1 | α1.1 (a) |
| igh_var19 | CIDRα1.1 (b) | B/A | EPCR | IGH | CIDRα1 | α1.1 (b) |
| raj116_var | CIDRα1.1 (c) | B/A | EPCR | raj116 | CIDRα1 | α1.1 (c) |
| ERS010178_NODE_17 | CIDRα1.8a | B/A | EPCR | GA026 | CIDRα1 | α1.8a |
| X2053_3 | CIDRα1.8b (a) | B/A | EPCR | GA027 | CIDRα1 | α1.8b (a) |
| ERS010532_NODE_326 | CIDRα1.8b (c) | B/A | EPCR | GA029 | CIDRα1 | α1.8b (c) |
| AMA1 | AMA1 | non-var | N/A | N/A | AMA1 | N/A |
| BSA | BSA | non-var | N/A | N/A | BSA | N/A |
| CSP | CSP | non-var | N/A | N/A | CSP | N/A |
| MSP1 | MSP1 | non-var | N/A | N/A | MSP1 | N/A |
| tetanus toxoid | tetanus toxoid | non-var | N/A | N/A | tetanus | N/A |

Listed antigens were used in a multiplex bead-based immunoassay to determine antigen-specific IgG reactivity of plasma from participants in the Kalifabougou chort. Bovine serum albumin (BSA) and tetanus toxoid were used as controls for non-specific cross-reactivity and the predictable response to tetanus vaccination, respectively.

**Figure S1. Study design.**
Participants and time points for used for "healthy baseline" and longitudinal analysis.

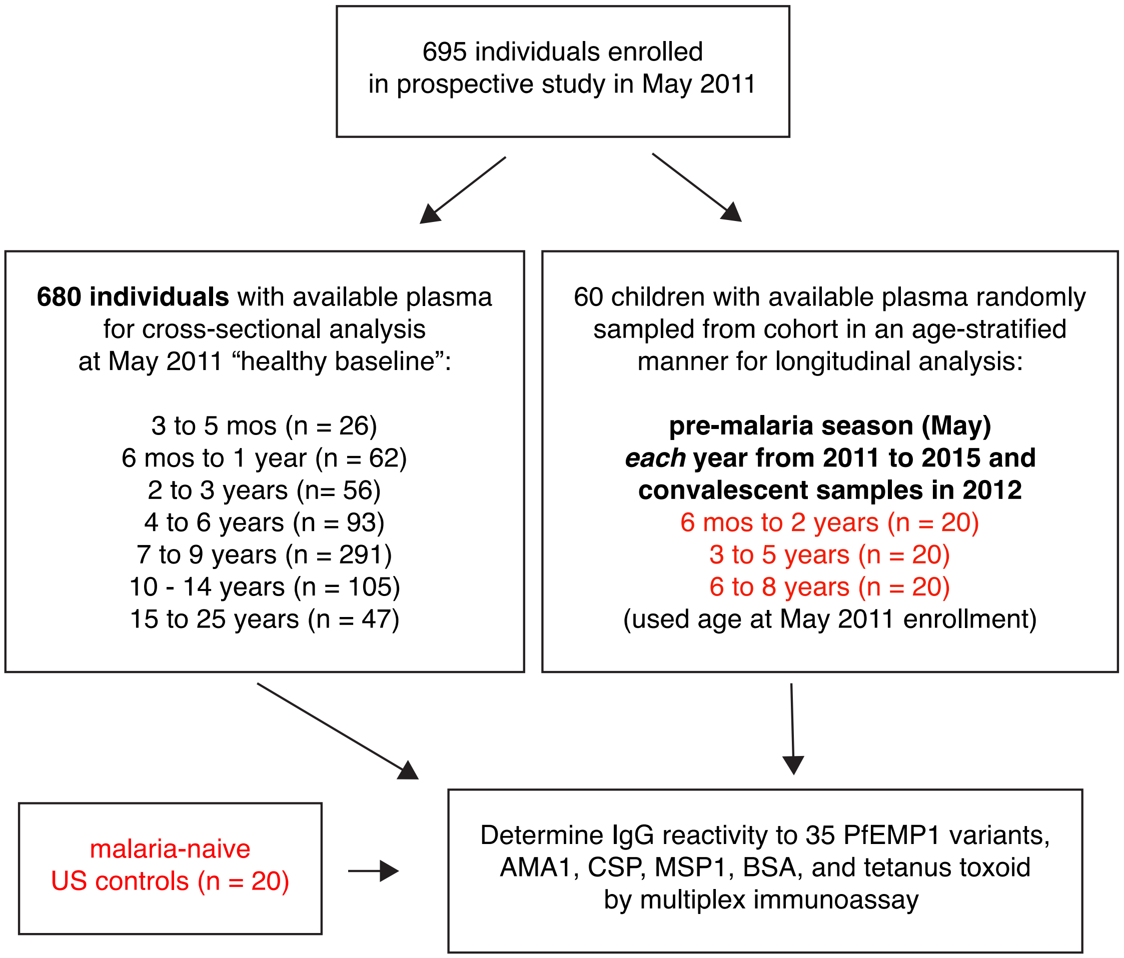

**Table S2. Differential acquisition of CIDR domain class-specific IgG antibodies with age and/or malaria exposure.**

| Domain Class | n | PfEMP1 group | Binding Phenotype | Slope for domain class | Coefficient (age:domain class interaction term) | Standard error | t | P value | BH-adjusted P value |
|---|---|---|---|---|---|---|---|---|---|
| CIDRγ | 340 | A | unknown | 0.224 | 0.0485 | 0.0134 | 3.63 | 0.000283 | 0.00141 |
| CIDRδ | 340 | A | unknown | 0.209 | 0.0357 | 0.00796 | 4.49 | <0.0001 | <0.0001 |
| CIDRα1 | 340 | A | EPCR | 0.208 | 0.0682 | 0.00435 | 15.7 | <0.0001 | <0.0001 |
| CIDRα2-6 | 340 | B | CD36 | 0.115 | -0.0935 | 0.00446 | -21.0 | <0.0001 | <0.0001 |

Refers to **Figure 1d**. To determine CIDR domain classes for which specific IgG was acquired more rapidly than the other variants, the change in variant-specific IgG reactivity with age was compared between all variants within each CIDR domain class and all other variants. Specifically, for each CIDR domain class, a linear regression model was performed for children <8 years of age, which represents the linear portion of the plot. The dependent variable was log-transformed antigen-specific IgG reactivity; the independent variables were presence of *P. falciparum* parasitemia (determined by PCR), age, and PfEMP-1 variant type dichotomized as the PfEMP-1 domain class of interest or variants in all other domain classes with the latter being the reference level. An interaction between age and CIDR class was included in the model. Tabulated coefficient and statistics are for the age:domain class interaction term. P value was adjusted for multiple testing (5 coefficients per model times 4 domain classes) using the Benjamini-Hochberg (BH) method. Non-PfEMP-1 antigens were not included in the analysis.

**Fig. S2. Antibodies specific for group A and EPCR-binding phenotypes are acquired earlier in life.**

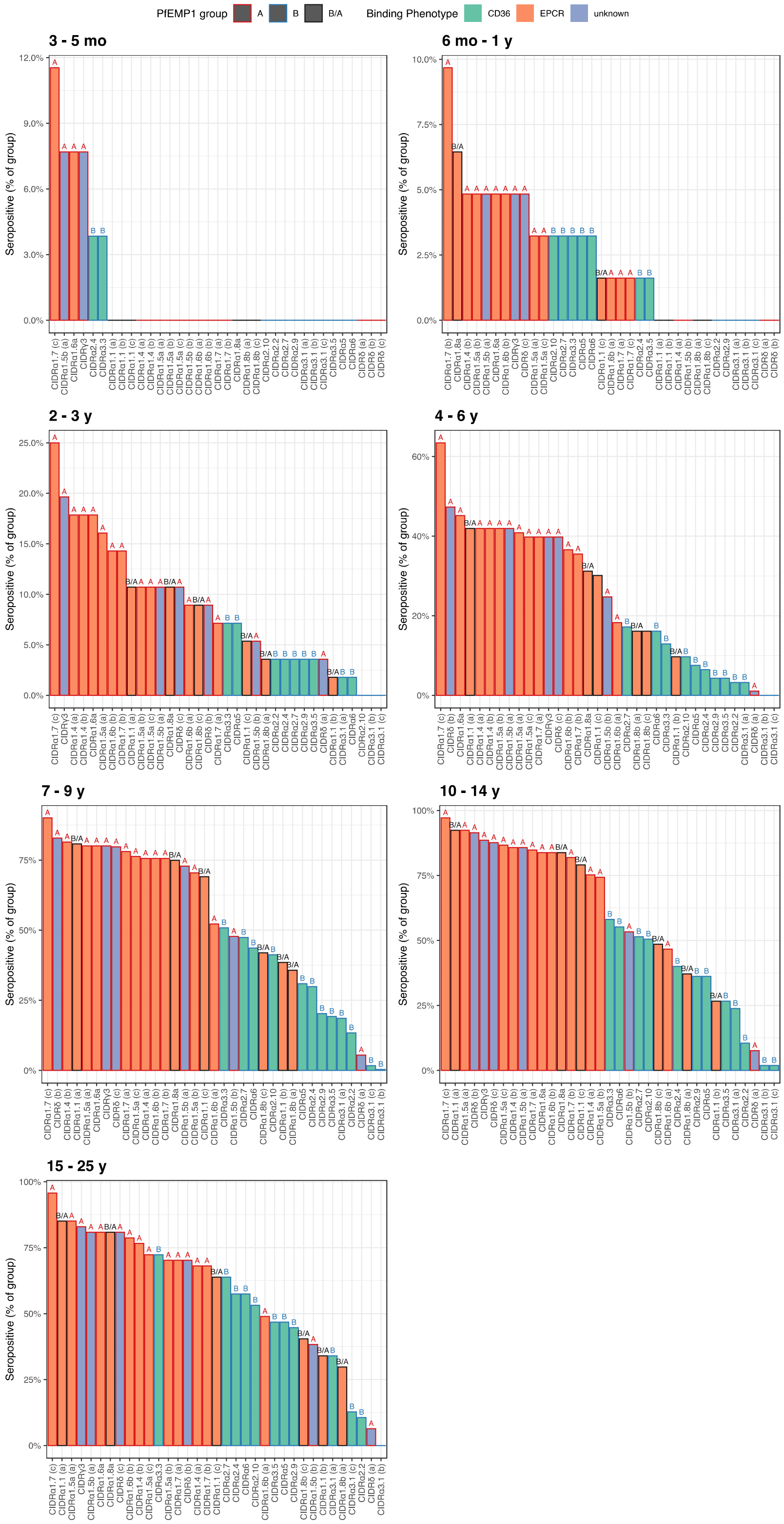


Antigens were ranked by seroprevalence to determine the dominant antibody responses for each age group. A response was considered seropositive if the antigen-specific IgG reactivity was greater than the mean reactivity of 20 malaria-naive US donors plus 3 standard deviations.

**Table S3. Relationship between CIDRγ3 seropositivity and protection from febrile malaria with inclusion of blood group O as a co-variate.**

| | without group O covariate | | | | with group O covariate | | | |
|---|---|---|---|---|---|---|---|---|
| **Covariate** | **HR** | **LCI** | **UCI** | **P value** | **HR** | **LCI** | **UCI** | **P value** |
| Age | 1.07 | 0.998 | 1.14 | 0.0582 | 1.07 | 0.999 | 1.14 | 0.055 |
| CIDRγ3 | 0.411 | 0.276 | 0.613 | 1.26E-05 | 0.407 | 0.274 | 0.605 | 8.73E-06 |
| group O blood type | | | | | 0.963 | 0.678 | 1.37 | 0.831 |
| Male | 0.858 | 0.605 | 1.22 | 0.39 | 0.859 | 0.606 | 1.22 | 0.395 |
| presence of HbS allele | 0.53 | 0.311 | 0.903 | 0.0196 | 0.529 | 0.311 | 0.902 | 0.0194 |

Results of Cox regression models assessing CIDRγ3-specific IgG on the risk of febrile malaria after incident *P. falciparum* infection in which covariates were age, gender, presence of the HbS allele, and AMA1-specific IgG reactivity without or with group O blood type. Analysis was restricted to children within the cohort who were at least 6 months of age, began the study negative for *P. falciparum* infection by PCR, and had ABO blood typing performed (218 subjects; 140 malaria events). Malaria risk was determined based on time to clinical malaria, defined as axillary temperature >37.5° C and any parasitemia, once parasitemia was detected by PCR. Results are ordered by increasing significance values.
